# Metagenomic unmapped reads provide important insights into human microbiota and disease associations

**DOI:** 10.1101/504829

**Authors:** Zifan Zhu, Jie Ren, Sonia Michail, Fengzhu Sun

## Abstract

We developed a computational pipeline, MicroPro, for metagenomic data analyses that take into account all the reads from known and unknown microbial organisms and for associating viruses with complex diseases. We utilized MicroPro to analyze metagenomics data related to three diseases: colorectal cancer, type-2 diabetes and liver cirrhosis, and showed that including reads from unknown organisms will markedly increase the prediction accuracy of the disease status based on metagenomics data. We identified new microbial organisms associated with these diseases. Viruses were shown to play important roles in colorectal cancer and liver cirrhosis, but not in type-2 diabetes. MicroPro is available at https://github.com/zifanzhu/MicroPro.

## Introduction

Trillions of microbes populate various sites of the human body and form microbiome communities [1]. These micro-organisms and their interactions between each other and the host play an important role in many physiological processes including metabolism, reproduction and immune system activity [2, 3]. In the nineteenth century, culture-based methods demonstrated that changes in these microbes might lead to disease. Since then, many subsequent studies confirmed these findings [4]. However, the cultivation technology only provided a limited view since many microorganisms could not be cultured in vitro [5]. Over the past twenty years, and thanks to the rapid development of sequencing technology, sequencing-based methods have gradually replaced the cultivation technology and have become the most widely used tools for microbial analysis. The 16S ribosomal RNA sequencing together with the recent shotgun whole genome sequencing not only discover large amounts of non-cultivable microbes, but also fundamentally change the way microbial analysis is performed [6, 7]. Researchers are now finding more evidence correlating human microbiota with various diseases such as colorectal cancer [8], type-2 diabetes [9, 10], liver cirrhosis [11] and many others. In addition, human microbiota has been linked to the effectiveness of cancer chemo-therapy [12]. In some studies, a single species or strain is associated with a disease while in other cases, groups of microorganisms interact to affect human health [13].

Mounting evidence connecting the microbiome with disease description has gradually brought about the concept of a supervised predictive study of microorganisms for different diseases. Although most of the studies are merely observational, which means we cannot simply conclude the causality between microbes and the disease [7], the existing correlations are sufficient to prove that performing a predictive study about the effect of microbiota on diseases is plausible. More specifically, many advances in this area have made it possible to predict the existence or states of a certain disease given information of the microorganisms for a specific subject.

In the field of machine learning, a supervised predictive study aims to build models based on sets of features to maximally approximate the response value or correctly classify the label of a sample. In the microbiota-disease setting, the response can either be disease/non-disease or different subtypes within a disease; thus a classification version of supervised predictive study is desired [14]. However, selection of features varies greatly among different studies. Our study is focused on analyzing the microbiome abundance in the context of shotgun whole genome sequencing. Similar analysis can also be applied to other choices of the feature including operational taxonomic units (OTUs, widely used in 16S rRNA analysis) [15], NCBI non-redundant Clusters of Orthologous Groups (COG) [16] or Kyoto Encyclopedia of Genes and Genomes (KEGG) groups [17]. With many software packages like MetaPhlAn2 [18] or Centrifuge [19] tackling the computation of the microorganisms’ abundance, the microbiota-disease predictive study can be formulated as a machine learning task based on a species-by-sample matrix with qualitative labels.

Recently, many studies have focused on the predictive analysis between human microbiota and diseases. For example, Zeller et al. [8] developed a species abundance based LASSO [20] model to differentiate between colorectal cancer patients and healthy indoviduals. Qin et al. [11] used gene markers to predict liver cirrhosis based on a Support Vector Machine (SVM) [21]. Moreover, Pasolli et al. [22] built a database named curatedMetagenomicData, which stored uniformly processed microbiome analysis results across 5,716 publicly available shotgun metagenomics samples. Using this database, Pasolli et al. developed a Random Forest [23] model to analyze the predictive power of different microbial features (such as species abundance, pathway coverage, etc.) on various diseases.

However, the current available approaches face a few challenges. Firstly, the feature characterization in microbiome studies often involves the process of mapping short reads against known microbial reference sequences in NCBI RefSeq database [24] or a catalogue of taxon-associated marker sequences [18]. However, a large proportion of the reads cannot be successfully mapped to a particular reference, which results in potential loss of valuable information. Secondly, the role of viruses in diseases is often neglected. Within human microbial community, bacteria reads constitute the majority while virus reads are reported as a very small proportion of the total reads (less than 5% in datasets used in our study). Additionally, incomplete database of viral reference genomes and the high mutation rates of viruses make them even more challenging to characterize and analyze [25]. Therefore, most disease-related microbiome studies focus only on the connection between bacteria and the disease. However, learning about the viruses is important as their abundance is about 10 times that of bacteria [26], and they can play important roles in multiple diseases. Norman et al. [27] showed that enteric virome change happened in patients with inflammatory bowel disease and bacteriophages might serve as antigens in human immune system. Ren et al. [28] demonstrated that decreased viral diversity was observed in patients with liver cirrhosis as compared to healthy individuals. Therefore, the role of viruses on human diseases should be investigated.

In order to overcome the challenges mentioned above, we developed a metagenomics predictive pipeline, MicroPro, which analyzes data in three main steps: 1). Reference-based known microbial abundance characterization: perform taxonomic profiling based on sequence alignment against reference genomes. 2). Assembly-binning-based unknown organism feature extraction: use cross-assembly to assemble the combined unmapped reads from all samples and consider that each assembled contig originates from an ‘unknown’ organism, which refers to an organism with no known reference available in database. Since some contigs may originate from the same organism, we cluster assembled contigs into bins and then treat each bin as an ‘unknown’ organism. 3). Machine learning predictive analysis: apply machine learning tools to predicting disease/non-disease or disease states based on the species-by-sample matrix. This is the first predictive pipeline based on a combination of both known and unknown microbial organisms. We tested MicroPro on four public NGS datasets and showed that consideration of unknown organisms markedly increased the prediction accuracy. Furthermore, we systematically investigated the effect of viruses on multiple diseases with the virus version of MicroPro. We examined the predictive power of the model with known and unknown viruses, and showed that unknown viruses played an important role in disease prediction warranting further attention.

## Results

### MicroPro: a metagenomics disease-related prediction analysis pipeline taking unmapped reads into consideration

We developed a new metagenomics analysis pipeline, MicroPro, to take into account both known and unknown microbial organisms for the prediction of disease status. MicroPro consists of three main steps: 1) Reference-based known microbial abundance characterization; 2) Assembly-binning-based unknown organism feature extraction; and 3) Machine learning predictive analysis. Figure 1 presents the procedures to extract the abundance table of both known and unknown microbial organisms. Various machine learning tools can then be applied to study the association between microbial abundances and the disease. Detailed explanations of each step are available in the Methods section.

**Figure 1.**
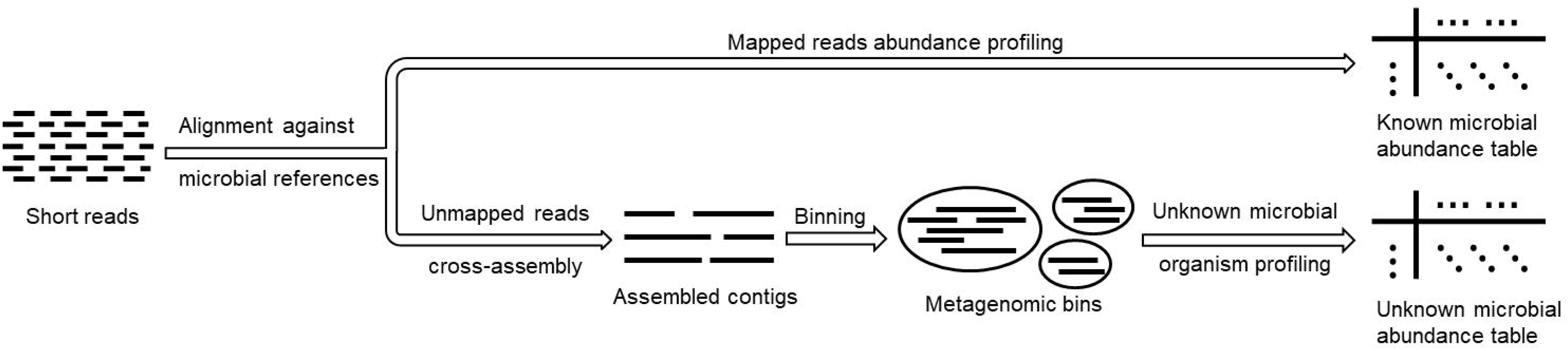
Feature extraction procedures of MicroPro.

We analyzed four publicly-available shotgun-sequenced metagenomics datasets related to three different diseases: Colorectal Cancer (CRC) [8], Type 2 Diabetes (T2D) [9, 10], and Liver Cirrhosis (LC) [11] (Table 1). These four datasets were all included in curatedMetagenomicData [22], a database containing various human microbiome features extracted from 5,716 publicly available shotgun metagenomics samples. Pasolli et al. [22] previously built a Random Forest model for each dataset based on known microbial species abundance extracted by MetaPhlAn2 [18]. We show that our prediction pipeline can give better prediction accuracy.

**Table 1:** Datasets analyzed in this paper.

| Dataset Name | Disease | Sample Size | # Cases | # Controls | Reference |
| --- | --- | --- | --- | --- | --- |
| ZellerG_CRC | CRC | 184 | 91 | 93 | Zeller et al. [8] |
| KarlssonFH_T2D | T2D | 96 | 53 | 43 | Karlsson et al.[9] |
| QinJ_T2D | T2D | 145 | 71 | 74 | Qin et al.[10] |
| QinN_LC | LC | 237 | 123 | 114 | Qin et al. [11] |

Detailed information of the four metagenomic datasets analyzed in this paper is provided. The dataset name is a combination of the author name and the corresponding disease.

### Adding abundance profile of unknown organisms markedly increased the predictive performance

The first step of MicroPro is to use Centrifuge [19] to characterize the relative abundance of known microbial species. Without adding unknown organism information, we directly trained a Random Forest classification model based on known microbial features. Its predictive power revealed the same level of accuracy as described by Pasolli et al. [22] which used MetaPhlAn2 [18] to characterize the relative abundance of known species. In general, the abundance characterization algorithm of Centrifuge and MetaPhlAn2 performed equally well in terms of prediction accuracy (Figure 2A).

**Figure 2.**
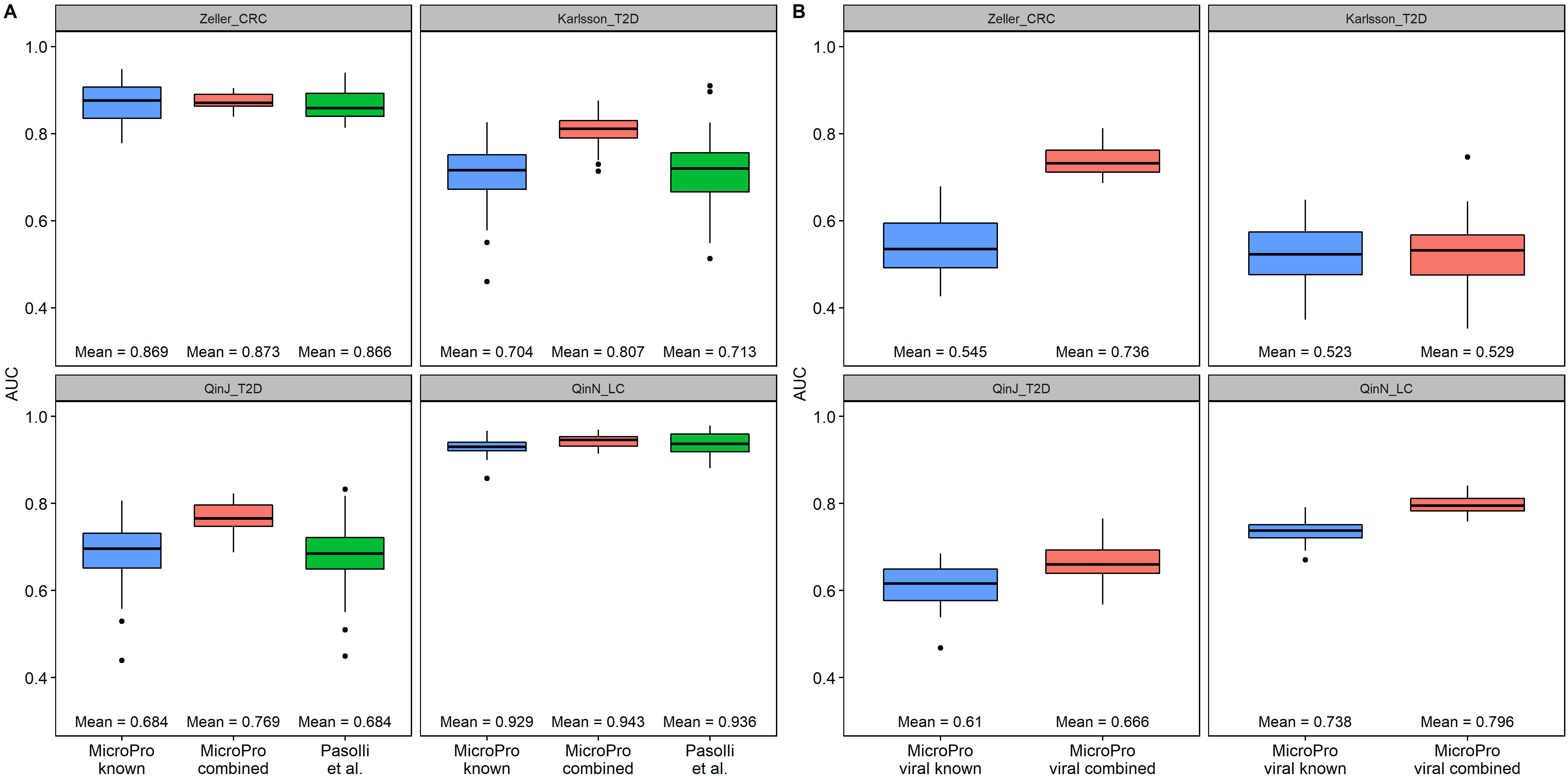
Prediction results of MicroPro. A). Boxplots of AUC scores obtained from using all the microbial features. “MicroPro known” refers to using only known microbial abundance profile extracted by MicroPro as the feature while “MicroPro combined” refers to using both known and unknown abundances. “Pasolli et al.” shows the prediction results of a similar study in [22], where microbial species abundance extracted by MetaPhlAn2 were used as the feature. Each model was repeatedly trained 30 times. Mean AUC scores are shown at the bottom of each plot. B). Boxplots of AUC scores obtained from only using viral features. “MicroPro viral known” refers to using only known viral abundance profile extracted by MicroPro while “MicroPro viral combined” refers to using both known and unknown viral abundances. Each model was repeatedly trained 30 times. Mean AUC scores are shown at the bottom of each plot.

The second step of MicroPro characterizes the abundance profiles from unknown organisms. We observed an obvious increase in the mean AUC scores when adding the abundances of unknown organisms. Also, all the four mean AUC scores are higher than the corresponding values in [22]. (Figure 2A) This result reveals the predictive value of the abundance profiles of unknown organisms that were commonly ignored by most of the currently available metagenomics analysis pipelines. It suggests that consideration of both known and unknown organisms significantly improves the prediction of metagenomics association analysis.

### Unknown viral abundances remarkably increased the AUC scores of virus-only prediction analysis

To test the predictive power of the viral organisms within the microbial community, we applied the virus version of MicroPro to all the four datasets. (Figure 2B) Although prediction accuracy using the abundance profiles of known viruses was much lower than the prediction accuracy using the abundance profiles of known microbials including bacteria, adding the unknown feature greatly improved the prediction accuracy for datasets ZellerG_CRC, QinN_LC and QinJ_T2D. This demonstrates the advantage of MicroPro to consider both known and unknown microbial organisms in metagenomics association study. On the other hand, we acknowledge that the increase in prediction accuracy for KarlssonFH_T2D is negligible. Considering the fact that there were only 28 unknown viral contigs found for this dataset, the abundance of unknown viruses were too small to play a role in the prediction analysis hence the low AUC increment. The relative low prediction accuracy for the two T2D datasets using both known and unknown viruses suggests that viruses may not play important roles in T2D. The relative high prediction accuracy for liver cirrhosis using known viruses suggests that some of the known viruses may play an important role in the disease. Similarly, the moderate increase of prediction accuracy for liver cirrhosis when unknown viruses are included suggests that some of the unknown viruses may also play some role in the disease. For colorectal cancer, the inclusion of unknown viruses significantly increased the prediction accuracy compared to cancer.

### Alpha diversity analysis of the abundance profiles of both microbial organisms and viruses

We also performed alpha diversity analysis for both microbial (Figure 3A) and viral (Figure 3B) abundance profiles in the cases and controls. Figure 3 shows the results of using the abundance profiles of both known and unknown microbial organisms. Alpha diversity results based on the abundance profiles of only known or unknown organisms are provided in Supplementary Figure 2 and 3, respectively. For microbial alpha diversity (Figure 3A), a consistent pattern of the case being less diverse is observed, which is consistent with the alpha diversity results for bacterial organisms in most studies. This pattern is most remarkable for QinN_LC, which corresponds to its high AUC score when using microbial abundances to differentiate between cases and controls (Figure 2A). We did not identify statistical significant differences for the viral alpha diversity between cases and controls for liver cirrhosis (QinN_LC) and type 2 diabetes (Karlsson_T2D, QinJ_T2D) at the type I error of 0.05. Surprisingly, we discovered that the viral diversity in CRC cases is much higher than that in the healthy controls, a finding consistent with the result from a recent study of Nakatsu et al. [29] who studied the viromes in CRC cases and controls.

**Figure 3.**
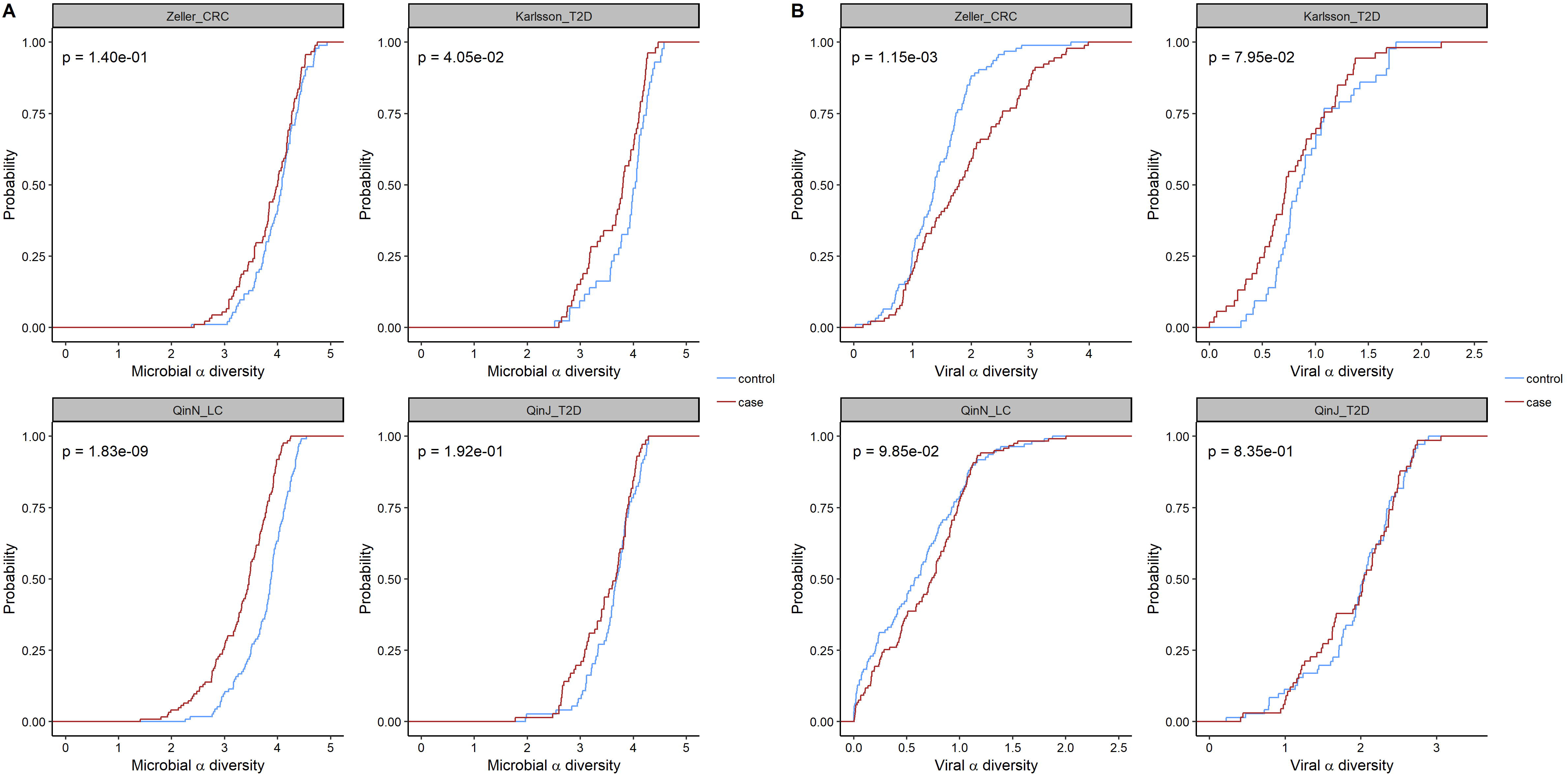
Cumulative probability of the alpha diversity. Plot A uses the abundance profiles of all the microbes while Plot B only uses the abundance profiles of viruses. For both plots, cumulative probability of alpha diversity with Shannon index is shown. Abundance profiles of both known and unknown organisms are used for the calculation. Pvalues based on the WMW test for the alpha diversity between the cases and the controls are provided.

### Significantly associated microbial organisms for each disease

We explored the microbial organisms that were significantly associated with a certain disease type in the metagenomics analysis. In our study, significantly associated microbial organisms were analyzed by investigating their importance scores in the Random Forest model. We repeatedly ran Random Forest model for 30 times and selected ten organisms with the highest importance scores. We considered an organism significant only if its selection frequency was above 18 in 30 runs. (Figure 4)

**Figure 4.**
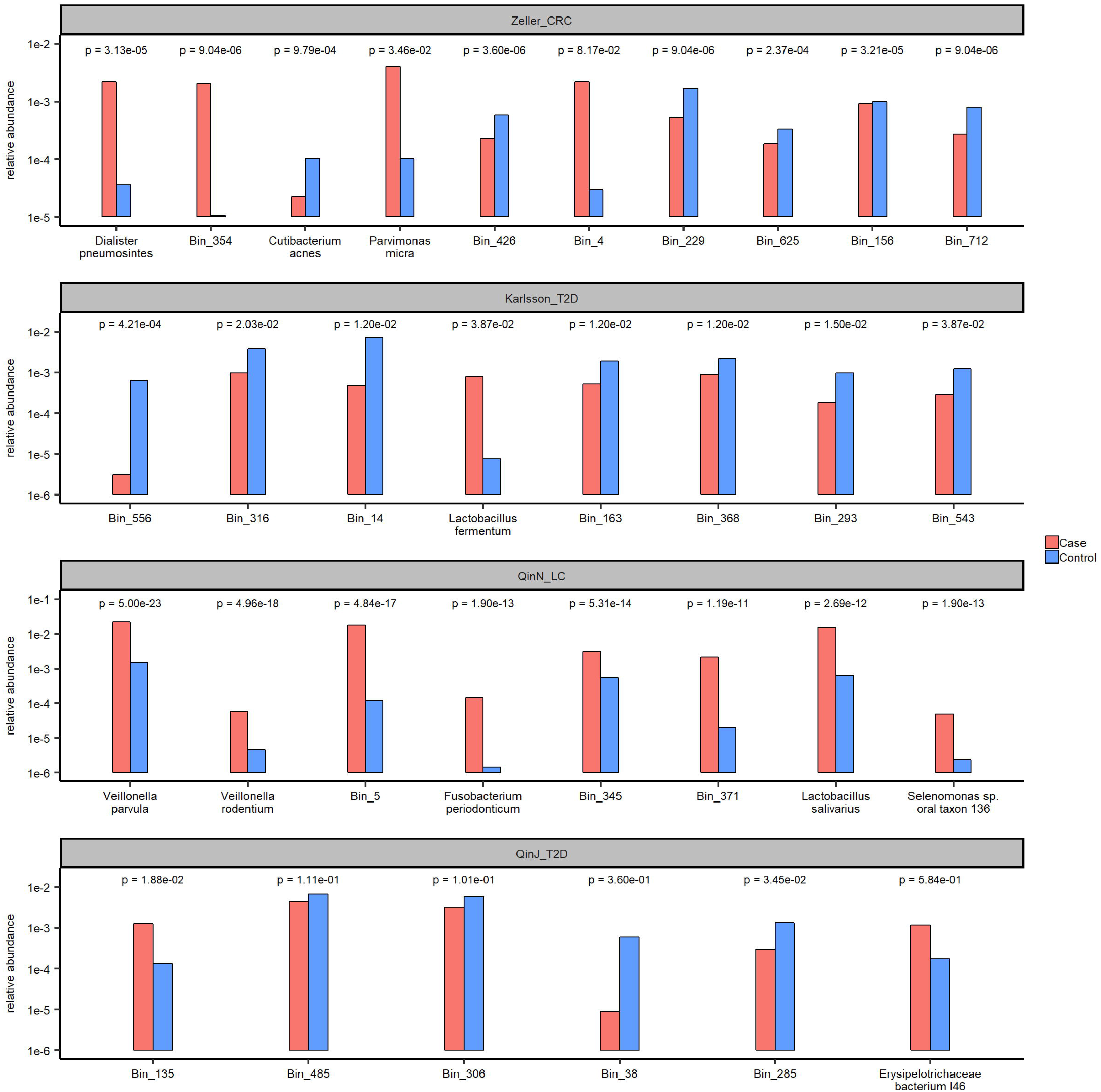
Significantly associated organisms for individual diseases. Each barplot shows the mean abundance of the organism in case and control. FDR adjusted WMW test p-values using the Benjamini-Hochberg procedure are provided. Known organisms are represented by their species name. Unknown organisms are named by “Bin_” together with a certain number. This number is merely for distinction purposes without biological relevance.

Some of our findings are consistent with results reported in the original studies. Karlsson et al. [9] showed that genus *Lactobacillus* was significantly more abundant in T2D patients than control. In Qin et al.’s work [11], *Veillonella parvula* and *Lactobacillus salivarius* were among the top 15 species that were more prevalent for liver cirrhosis patients. However, due to the tremendous increase in the reference genome database sequences, we found several new species associated with diseases that were not included in the database of original papers.

For example, we discovered a number of organisms which were predictive of liver cirrhosis (Figure 4). These organisms include *Veillonella parvula, Veillonella rodentium, Fusobacterium periodonticum, Lactobacillus salivarius* and *Selenomonas sp. oral taxon 136*. These organisms frequently inhabit the oral cavity and many are pathogenic. For example, *Veillonella parvula* is a bacterium in the genus *Veillonella. Veillonella* are Gram-negative bacteria anaerobic cocci. *Veillonella parvula* is well known for its lactate fermenting abilities and inhabit the intestines and oral mucosa. In humans, *Veillonella* can cause osteomyelitis, endocarditis, periodontitis and dental caries as well as various systemic infections [30]. Similarly, *Fusobacterium* is a genus of anaerobic, Gram-negative, non-spore-forming bacteria, similar to *Bacteroides*. Although in the past, *Fusobacterium* was considered part of the normal oral microbiome, the current consensus is that *Fusobacterium* should always be treated as a pathogen [31] and has been linked to periodontal diseases, ulcerative colitis, and colon cancer. These organisms originate from the mouth, but may also inhabit the intestine [32]. Even though our model discovered new organism-associations for disease prediction, it has been shown that the oral microbiota can influence the gut microbiome and has been detected in the stools of patients with cirrhosis [11]. Chen et al. [33] described *Veillonella* and other oral microbiota as discriminative taxa between patients with cirrhosis compared to controls. The permissive oral microbial invasion may be related to altered hepatic bile production or the frequent use of proton pump inhibitors in this population. Both bile and gastric acid are natural gates that can inhibit the survival of many of the ingested organisms. Furthermore, bacterial populations originating from the oral microbiota are capable of producing high levels of methyl mercaptan (CH3SH). Elevated blood levels of CH3SH have been linked to the development of hepatic encephalopathy [34].

The presence of both *Dialister pneumosintes* and *Parvimonas micra* was predictive of development of colorectal cancer in our model. *Dialister pneumosintes* was found in patients with periodontitis [35] and has been shown to have potential pathogenic roles in various human body sites including the lung and brain [36]. It has been recently shown to be an important component of the dysbiotic microbiome in patients with gastric cancer [37]. *Parvimonas micra* can cause infectious endocarditis [38], native joint septic arthritis [39] and spondylodiscitis [40] and has also been associated with gastric cancer [37]. Not only enrichment of specific organism was predictive of colorectal cancer in our model, but we also report depletion of specific organisms, such as *Cutibacterium acnes*, is seen in association with this type of cancer. While this organism was originally described in subjects with acne, it can still be found throughout the digestive tract [41] and was originally named *Propionibacterium acnes* for its ability to generate propionic acid [42]. Propionic acid, among other short chain fatty acids (SCFA), contributes to the health of colonocytes and has been shown to be depleted in colorectal cancer [43]. The discovery that subjects with colorectal cancer harbor less cutibacterium acnes could potentially explain the previous reports of depletion of propionic acid in this population and may shed some light on the pathophysiology of disease development.

Another interesting observation was the association of some unknown organisms with the corresponding diseases. We have demonstrated that adding the abundance profile of unknown organisms would remarkably increase the prediction accuracy. Since unknown organisms made up a large proportion of the organisms that are disease-associated, it was not surprising to see that many were predictive of disease, further highlighting the advantage of our pipeline since otherwise these unknown features would not be explored.

### Prediction analysis between two T2D datasets

Although prediction within one study can usually give good results, prediction accuracy drops sharply when applied to a different dataset. Different experiment protocols, various sequencing platforms, variable time points of data collection are all possible reasons for the drop in prediction accuracy. In our study, there were two T2D datasets, which allowed the determination of the generalization potential of the predictive model across different studies. As shown in Figure 5, the AUC scores dropped markedly for both cases when compared with prediction within one study (Figure 2A). When using KarlssonFH_T2D to predict QinJ_T2D, adding the unknown feature seemed to have no effect on the prediction accuracy. However, in the other case, adding the unknown features helped to increase the AUC from 0.61 to 0.64 suggesting that in cross-study settings, adding unknown organisms can result in higher prediction accuracy.

**Figure 5.**
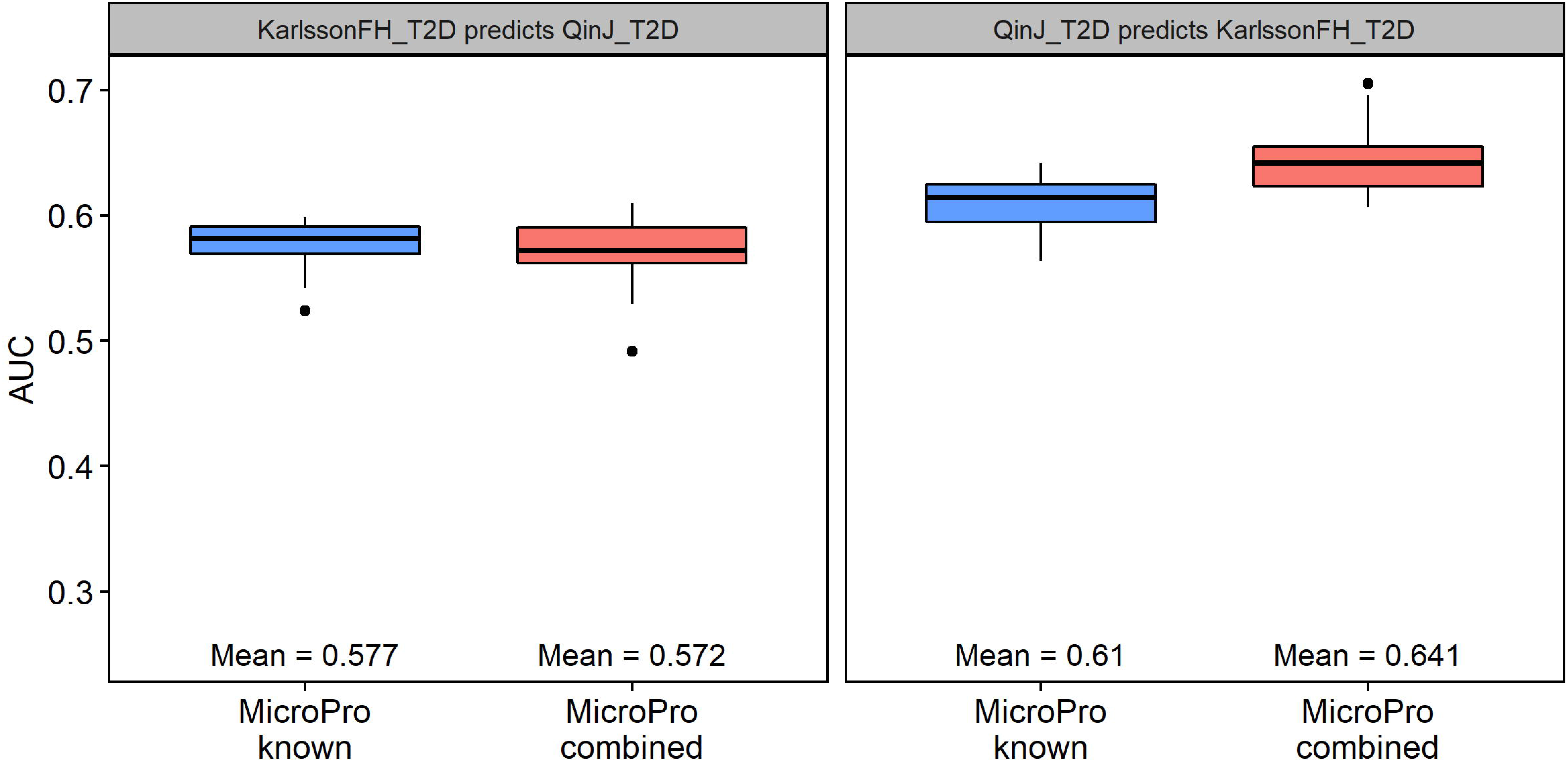
Prediction analysis between two T2D datasets. “MicroPro known” refers to using only known microbial abundance profile extracted by MicroPro as the feature while “MicroPro combined” refers to using both known and unknown abundances. Each model was repeatedly trained 30 times. Mean AUC scores are shown at the bottom of each plot

## Discussion

Many studies have described the development of computational tools to investigate the association of microbial organisms with complex traits. However, most of the available tools focus on the microbial species with a known reference genome and the reads not mapped to the known genomes are not considered, which can result in loss of potentially useful information. In order to address this issue, we developed the MicroPro pipeline that extracts both known and unknown microbial features within metagenomic datasets. We tested MicroPro in a disease prediction study involving four public metagenomic datasets covering three different diseases. We show that the prediction accuracy is markedly increased when adding unknown microbial features, which demonstrates the important predictive role of unknown organisms that may have been overlooked by most metagenomic studies.

Many studies have demonstrated the important role of viruses in human diseases like inflammatory bowel disease [27] and liver cirrhosis [28]. However, due to the limited virus genome database and high mutation rates, viruses were often neglected in metagenomic association studies. The virus version of MicroPro aims at extracting both known and unknown viral features from sequenced reads. We performed prediction analysis with viral abundances extracted by the virus version of MicroPro on the same public metagenomic datasets. The results indicated that viruses did play some roles in diseases like colorectal cancer and liver cirrhosis. Thus, the role of viruses should not be ignored in metagenomic analysis. Also, for some datasets, like ZellerG_CRC in our study, the power of predicting disease when using known virus only, was no better than random guess. However, the inclusion of unknown viral features remarkably increased the prediction accuracy. Therefore, our pipeline was able to distinguish the role of viruses by investigating unknown features.

We also discovered many novel microbial associations with specific diseases and disease prediction. Some of these associations are consistent with what has been described in the past. Even though we discovered new organisms predicting liver cirrhosis, however, those germs seem to originate from the mouth, a concept that is consistent with prior findings. Nevertheless, these discoveries may set the foundation for development of specific biomarkers of disease, which may be specific to the novel organisms identified in this study. Other discoveries were unique, such as identifying the depletion of *Cutibacterium acnes*, in colorectal cancer. *Cutibacterium acnes* generates propionic acid, which is one of the SCFA contributing to the health of the colonocytes and is known to be depleted in colorectal cancer [43]. Such association can lay ground work for future mechanistic understanding of the pathophysiology of this disease.

We acknowledge that there are limitations of our pipeline. Although unknown features are extracted through assembly and binning, more functional analysis is needed to further understand the roles each bin playing in diseases. Also, the disease prediction study is only observational and does not show the causality between a certain or a group of microbes and diseases. Furthermore, though we only tested MicroPro in disease related analysis, MicroPro is readily to be applied to any type of phenotype prediction metagenomics studies. By fully utilizing both known and unknown organisms including viruses in microbiota, we expect MicroPro will help to largely improve the prediction accuracy and facilitate biomarker detections.

## Conclusions

MicroPro provides a highly useful tool to study the associations between microbiota and diseases without neglecting key information from unknown organisms. The microbial prediction of disease can be useful in understanding disease pathogenesis and may become crucial in laying ground work for future development of specific disease biomarkers.

## Methods

### Datasets

We downloaded all the datasets using the links provided in the original papers [8–11]. We denote each data by the last name, followed by the initial of the given name of the first author and then the disease. The number of cases and controls are given in Table 1. For ZellerG_CRC, the “small adenoma” samples were treated as controls while the “large adenoma” samples were removed.

Cross-assembly needed large amounts of computing resources. We successfully cross-assembled the ZellerG_CRC, KarlssonFH_T2D and QinJ_T2D without losing any data. However, for the largest dataset QinN_LC, after using all its reads for known microbial abundance characterization, we performed a downsampling and used 70% (which was the best we could do) of the unmapped reads to do cross-assembly due to limited computing memories.

### MicroPro: a pipeline of predicting phenotypes based on metagenomics data

#### Step 1: Reference-based known microbial abundance characterization

We used Centrifuge [19] to map the reads to the microbial genomes and calculated the abundance profiles of known microbial organisms from the metagenomics data. Centrifuge is an alignment-based taxonomic profiling tool. Its microbial database contains all the available bacterial, viral, and archaeal reference genomes in NCBI, updated on Jan 4, 2018. Centrifuge also utilizes an Expectation-Maximization (EM) algorithm to compute the abundance for each microbial species. This EM based algorithm is similar in spirit as those used in Cufflinks [44], Sailfish [45] and GRAMMy [46]. It performs well especially in the presence of reads mapped to multiple genomes and multiple locations in the same genome. In our study, we adopted the species abundance calculated by Centrifuge as the known microbial feature.

#### Step 2: Estimating abundance profiles of unknown microbial organisms based on reads assembly followed by contig binning

Although Centrifuge accurately characterizes known microbial relative abundance profiles, a large fraction of reads cannot be mapped to the known microbial organisms. The average mapping rate for each dataset is about 35% - 40% in our study (Supplementary Figure 1). The large amount of unmapped reads can potentially provide extra information on the prediction accuracy of phenotypes based on the metagenomics data. Therefore, our main objective in this step is to take into account the unmapped reads for phenotype prediction.

After filtering out mapped reads from the metagenomics data, we performed cross-assembly on all samples using Megahit [47]. Megahit efficiently assembles large and complex metagenomics data *de novo* based on succinct de Bruijin graph. After cross-assembly, we used MetaBAT 2.12.1 [48] to perform binning on the assembled contig set. MetaBAT 2.12.1 is a reference-free metagenomics binner and its binning criterion is based on tetranucleotide frequency and mean base coverage. This “reference-free” feature is crucial to our study, since the contig set to be binned contained no reads that could be mapped to a known reference. Recent comparative studies on contig binning [49] showed that MetaBAT 2.12.1 performs well compared to other algorithms for conting binning.

Reads assembly and contig binning are highly important to recover unknown organisms from the unmapped reads. Here, ‘unknown organisms’ represents the organisms without a known reference. Once we finished cross-assembly and metagenomics binning, we treated each contig bin as an unknown organism and the binned reads as a part of its genome. In terms of defining the feature of the unknown organisms, we still used the abundance, just as what we did for known species. The formula of the abundance (Ab) of unknown organism *i* was:

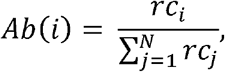

where *rc* was the length normalized read counts, which was defined as the number of reads mapped to that organism divided by its genome length. Here calculating *rc* was a major issue, since we do not know the whole genome of the unknown organism. To overcome this challenge, we first mapped all the unmapped reads back to the contig set using BWA-aln [50] with parameter “-n” set as 0.03 (only alignments with more than 97% accuracy were considered mapped). Then we calculated the length normalized read counts (*rc*) for each contig according to the mapping results. Finally, for each contig bin (i.e each unknown organism), we took the average *rc* of all the contigs that belonged to it as an approximation of its real *rc*. Thus, we could compute the unknown feature for all contig bins using the above formula. The next step was to combine the known feature and the unknown feature, and perform the prediction analysis.

#### Step 3: Predicting phenotypes using Random Forests

In the above two steps, we extracted the relative abundance profiles of both known and unknown microbial organisms. Then, it is natural to combine them and train a machine learning model to differentiate between the cases and the controls. The machine learning tool we chose in our study was Random Forests [23]. It is an ensemble of the decision tree algorithm and is highly robust to over-fitting when the number of features is far greater than the number of samples. We randomly separated the dataset into training set and test set with ratio 7:3. During model training, we adopted 10-fold cross-validation to tune the parameters for best predictive performance. In terms of the measure of prediction accuracy, we adopted the Area Under the receiver operating characteristic Curve (AUC) score, a widely used performance measure of the classification model. An AUC score close to 1 indicated perfect classification, while a 0.5 AUC score revealed that the model was as poor as a random guess. We repeated the above procedure for 30 times and the average AUC score was reported.

Pasolli et al. [22] built a Random Forest model for all four datasets based on known microbial abundance extracted by MetaPhlAn2 [18]. However, they trained a model on the whole dataset and tested it back to the data itself. This procedure would under-estimate the true test error since the test data was involved in model training. We modified the R-script provided by the author to allow train-test splitting for a fair comparison and obtained the AUC scores presented in Table 1.

### Predicting phenotypes based on virus abundance profiles

Viruses play a very important role in human microbial community by controlling the balance of different bacterial organisms. However, due to its relative low abundance, extraction of all the viral information, especially those without a known reference, remains a major difficulty. Aimed at making full use of all the viral features within metagenomics samples, the virus version of MicroPro is similar in spirit to the general Pipeline presented in the previous section, except that we add a step of viral contigs detection. The full pipeline is shown below.

#### Step 1: Known viral abundance extraction

For the known viral abundance, we still used the software Centrifuge. But this time, we only included the viral reference genomes as our database. Centrifuge mapped metagenomics samples against all the viral genomes, and then calculated their abundance based on an EM algorithm. Again, we adopted the abundance in the Centrifuge output as the known viral feature.

#### Step 2: Unknown viral feature detection

We performed cross-assembly on the unmapped reads filtered out by Centrifuge results. Before metagenomic binning, we applied VirFinder [28] for viral contigs detection. VirFinder utilized a logistic regression model to differentiate between bacteria and virus contigs. We considered a contig as virus if its VirFinder q-value is smaller than 0.2. Q-vaule [51] is a p-value correction method targeting exact false discovery rate (FDR) control. We performed metagenomic binning on the viral contigs and calculated viral bins’ abundance using the same method as described in the previous section Step 2.

#### Step 3: Predicting phenotypes based on viral abundance

With both the known and unknown viral features at hand, the next step was to perform the prediction analysis. We combined two viral features and trained a Random Forest model based on extracted viral abundance. We used 10-fold cross-validation to tune the parameters and set AUC score as the measure of prediction accuracy.

### Alpha diversity analysis

Alpha diversity is a widely-used diversity measure in microbiome studies It is defined based on both the number of species within a sample and the abundance of each species. We performed alpha diversity analysis of both microbial and viral abundance profiles. Alpha diversity with Shannon index is calculated by package “vegan” in R.

### Significantly associated microbial organisms for each disease

We found out the significantly associated features and investigate their roles by comparing the importance scores, which was a commonly-used criterion for the Random Forest (RF) model. A variable with a higher importance score meant that it has a higher splitting value in the model. For each dataset, we repeatedly trained the RF model for 30 times, with both known and unknown microbial abundance included in the feature. During each run, we sorted the variables by importance scores and selected the top 10 of them. After 30 runs, we counted the selection frequency of each variable, and calculated their average importance scores. Note that importance score ranged from 0 to 100.

### Predictive study between the two T2D datasets

We trained a Random Forest model based on one of the T2D datasets and tested on the other to obtain the AUC score. Features included were also the known and unknown microbial abundance. Obtaining the known feature was essentially the same procedure as MicroPro’s Step 1. We used the following strategy to calculate the abundance profiles of the unknown microbial organisms. For the train set, we used MicroPro’s Step 2 to find out the unknown microbial feature. For the test set, instead of mapping back to its own contig set, we aligned the unmapped reads in the test set against the train data contig set. In this way, we could obtain a consistent feature matrix so that the following prediction analysis could be performed seamlessly.

## Declarations

### Ethics approval and consent to participate

Not applicable.

### Consent for publication

All authors have approved the manuscript for submission.

### Availability of data and materials

All the datasets used in this study are publicly available from the European Nucleotide Archive (ENA) database (http://www.ebi.ac.uk/ena). Accession number for ZellerG_CRC is ERP005534 [8], for KarlssonFH_T2D is ERP002469 [9], for QinN_LC is ERP005860 [11], for QinJ_T2D is SRA045646 [10].

### Competing interests

The authors declare that the research was conducted in the absence of any commercial or financial relationships that could be construed as a competing interest.

### Funding

National Science Foundation (NSF) [DMS-1518001]; National Institutes of Health (NIH) [R01GM120624].

### Authors’ contributions

JR and FS conceived and designed the study. JR, ZZ and FS designed the methodological framework. ZZ implemented the methods, carried out the computational analyses and drafted the paper. SM explained the possible roles of the significantly associated microbial organisms found in the analysis. JR and SM modified the paper. SM and FS finalized the paper. All authors agree to the content of the final paper.

## Acknowledgements

This research utilized resources of HPC (High-Performance Computing), which is supported by the University of Southern California.

## Additional files

**Additional Figure 1.**
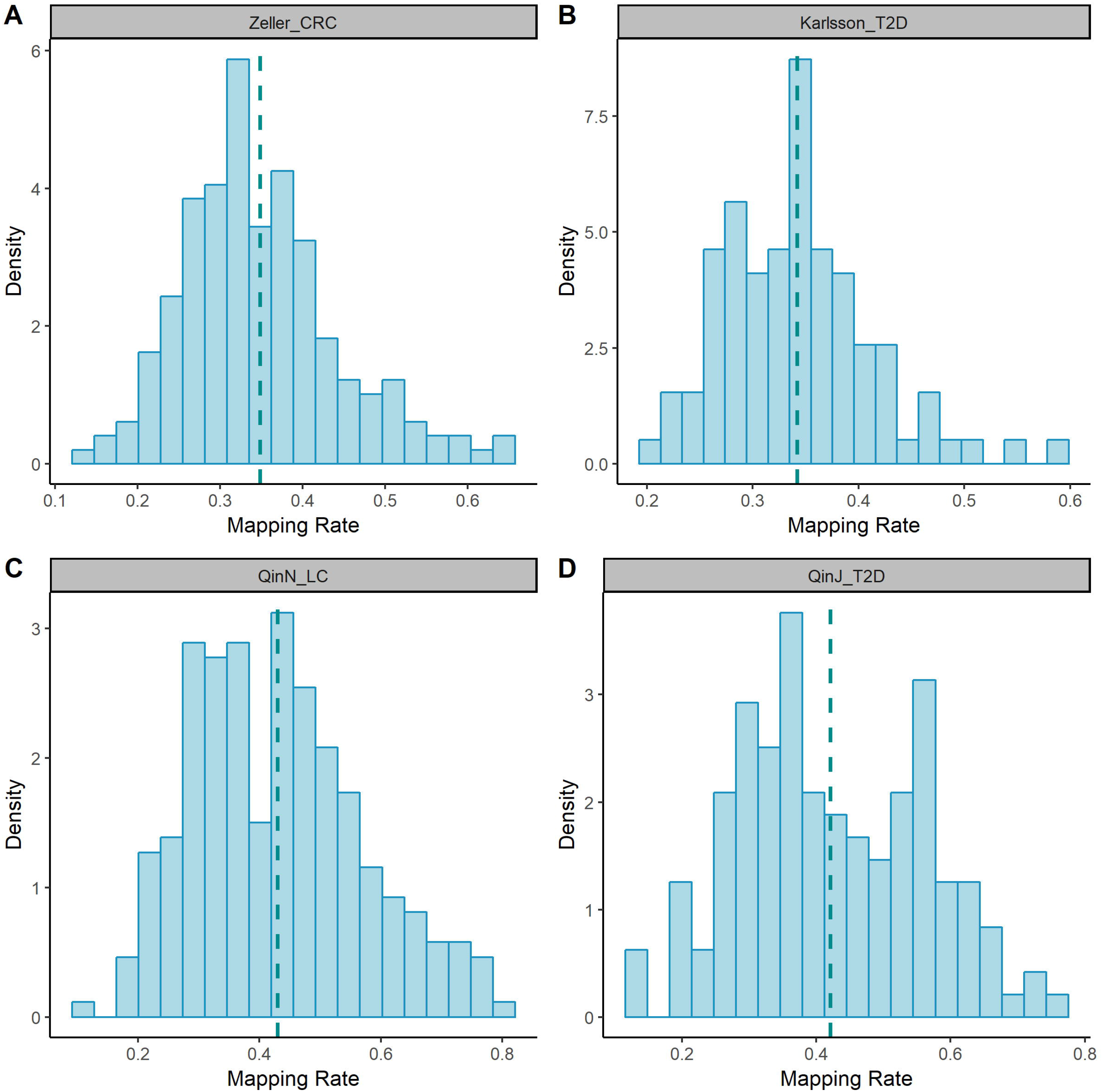
Histograms of mapping rates of each dataset. Dashed lines show the mean mapping rate.

**Additional Figure 2.**
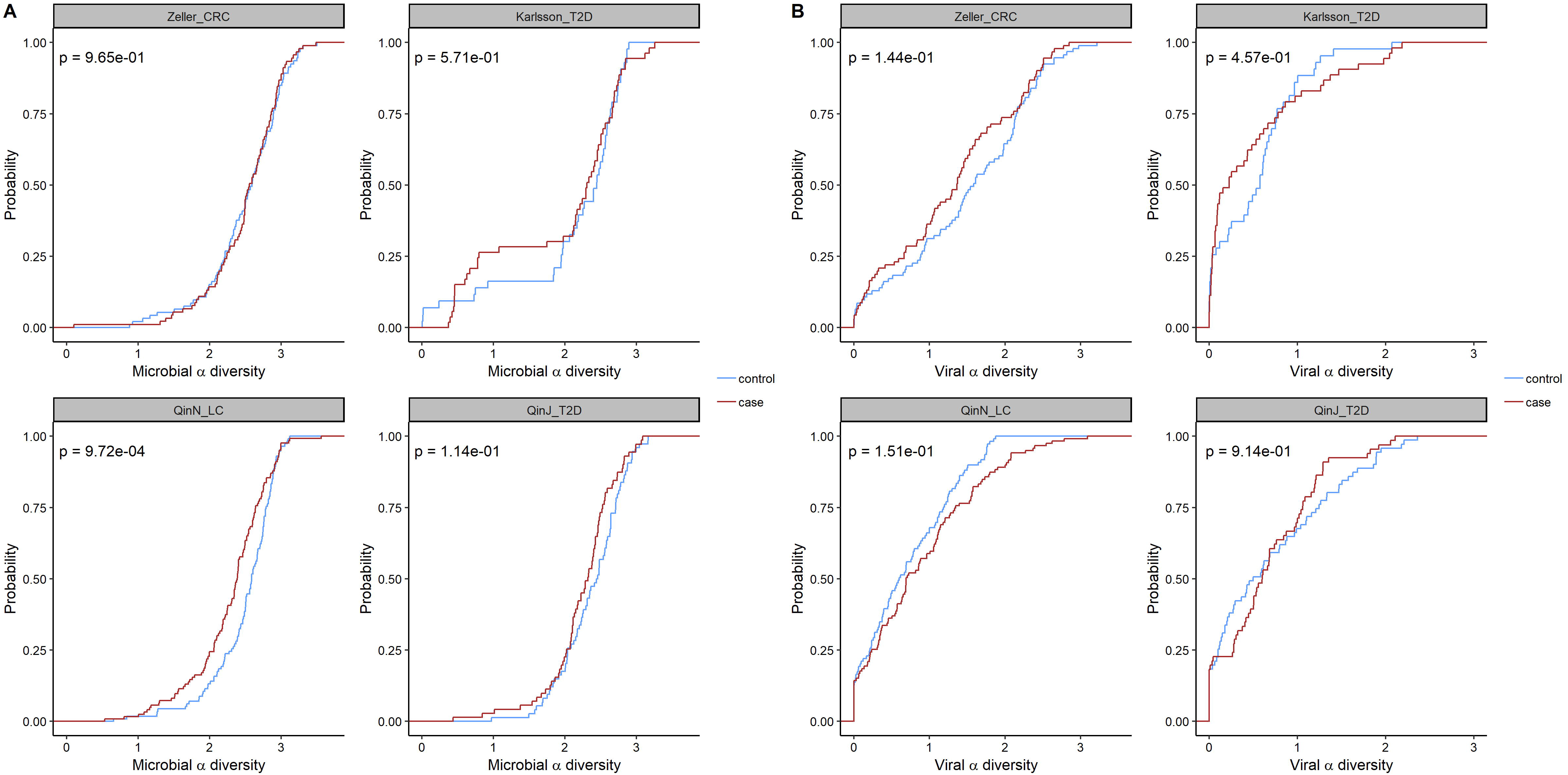
Cumulative probability of alpha diversity of known profile. Plot A uses all the microbial abundances while Plot B only uses viral abundances. For both plots, only known abundances are used for the calculation. Shannon index is set as the diversity index. WMW test pvalues between the cases and the controls are provided.

**Additional Figure 3.**
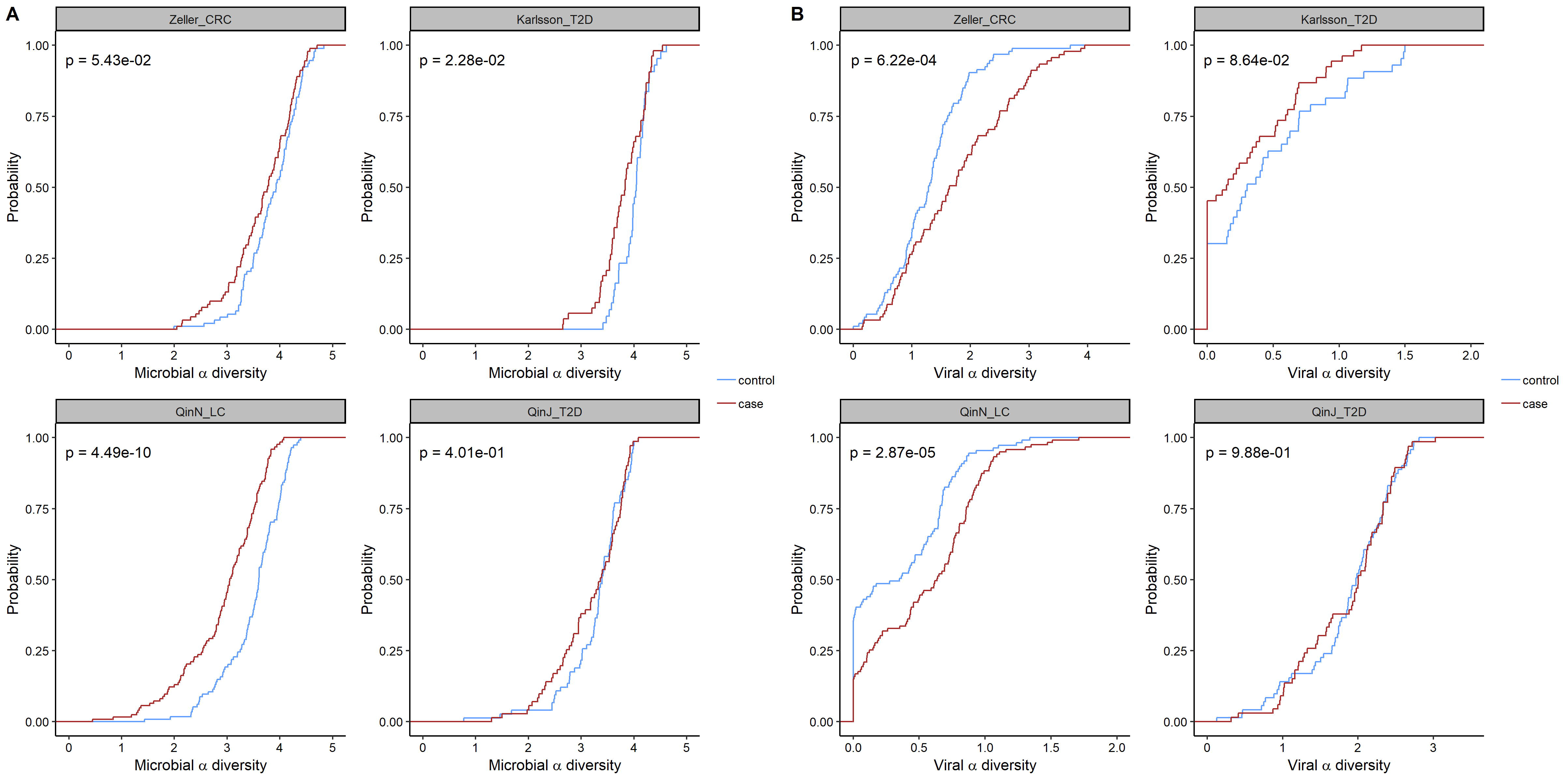
Cumulative probability of alpha diversity of unknown profile. Plot A uses all the microbial abundances while Plot B only uses viral abundances. For both plots, only unknown abundances are used for the calculation. Shannon index is set as the diversity index. WMW test pvalues between the cases and the controls are provided.

## References

1. Savage, D.C., Microbial ecology of the gastrointestinal tract. Annu Rev Microbiol, 1977. 31: p. 107–33.

2. Mazmanian, S.K., et al., An Immunomodulatory Molecule of Symbiotic Bacteria Directs Maturation of the Host Immune System. Cell, 2005. 122(1): p. 107–118.

3. Benson, A.K., et al., Individuality in gut microbiota composition is a complex polygenic trait shaped by multiple environmental and host genetic factors. Proceedings of the National Academy of Sciences, 2010. 107(44): p. 18933.

4. Dethlefsen, L., M. McFall-Ngai, and D.A. Relman, An ecological and evolutionary perspective on human-microbe mutualism and disease. Nature, 2007. 449(7164): p. 811.

5. Baumann, P. and N.A. Moran, Non-cultivable microorganisms from symbiotic associations of insects and other hosts. Antonie van Leeuwenhoek, 1997. 72(1): p. 39–48.

6. Tannock, G.W. What immunologists should know about bacterial communities of the human bowel. in Seminars in immunology. 2007. Elsevier.

7. Cho, I. and M.J. Blaser, The human microbiome: at the interface of health and disease. Nature Reviews Genetics, 2012. 13(4): p. 260.

8. Zeller, G., et al., Potential of fecal microbiota for early-stage detection of colorectal cancer. Molecular systems biology, 2014. 10(11): p. 766.

9. Karlsson, F.H., et al., Gut metagenome in European women with normal, impaired and diabetic glucose control. Nature, 2013. 498(7452): p. 99.

10. Qin, J., et al., A metagenome-wide association study of gut microbiota in type 2 diabetes. Nature, 2012. 490(7418): p. 55.

11. Qin, N., et al., Alterations of the human gut microbiome in liver cirrhosis. Nature, 2014. 513(7516): p. 59.

12. Zitvogel, L., et al., The microbiome in cancer immunotherapy: Diagnostic tools and therapeutic strategies. Science, 2018. 359(6382): p. 1366–1370.

13. Knights, D., et al., Human-associated microbial signatures: examining their predictive value. Cell host & microbe, 2011. 10(4): p. 292–296.

14. Knights, D., E.K. Costello, and R. Knight, Supervised classification of human microbiota. FEMS microbiology reviews, 2011. 35(2): p. 343–359.

15. Schloss, P.D. and J. Handelsman, Introducing DOTUR, a computer program for defining operational taxonomic units and estimating species richness. Applied and environmental microbiology, 2005. 71(3): p. 1501–1506.

16. Von Mering, C., et al., STRING: known and predicted protein-protein associations, integrated and transferred across organisms. Nucleic acids research, 2005. 33(suppl_1): p. D433–D437.

17. Kanehisa, M., et al., The KEGG resource for deciphering the genome. Nucleic acids research, 2004. 32(suppl_1): p. D277–D280.

18. Truong, D.T., et al., MetaPhlAn2 for enhanced metagenomic taxonomic profiling. Nature methods, 2015. 12(10): p. 902.

19. Kim, D., et al., Centrifuge: rapid and sensitive classification of metagenomic sequences. Genome research, 2016.

20. Tibshirani, R., Regression shrinkage and selection via the lasso. Journal of the Royal Statistical Society. Series B (Methodological), 1996: p. 267–288.

21. Hearst, M.A., et al., Support vector machines. IEEE Intelligent Systems and their applications, 1998. 13(4): p. 18–28.

22. Pasolli, E., et al., Accessible, curated metagenomic data through ExperimentHub. Nature methods, 2017. 14(11): p. 1023.

23. Breiman, L., Random forests. Machine learning, 2001. 45(1): p. 5–32.

24. Pruitt, K.D., et al., RefSeq: an update on mammalian reference sequences. Nucleic acids research, 2013. 42(D1): p. D756–D763.

25. Bonhoeffer, S. and P. Sniegowski, Virus evolution: the importance of being erroneous. Nature, 2002. 420(6914): p. 367.

26. Breitbart, M. and F. Rohwer, Here a virus, there a virus, everywhere the same virus? Trends in microbiology, 2005. 13(6): p. 278–284.

27. Norman, J.M., et al., Disease-specific alterations in the enteric virome in inflammatory bowel disease. Cell, 2015. 160(3): p. 447–460.

28. Ren, J., et al., VirFinder: a novel k-mer based tool for identifying viral sequences from assembledmetagenomic data. Microbiome, 2017. 5(1): p. 69.

29. Nakatsu, G., et al., Alterations in Enteric Virome Associate With Colorectal Cancer and Survival Outcomes. Gastroenterology, 2018.

30. Mashima, I., et al., Exploring the salivary microbiome of children stratified by the oral hygiene index. PloS one, 2017. 12(9): p. e0185274.

31. Aliyu, S.H., et al., Real-time PCR investigation into the importance of Fusobacterium necrophorum as a cause of acute pharyngitis in general practice. Journal of medical microbiology, 2004. 53(10): p. 1029–1035.

32. Zoetendal, E.G., et al., The human small intestinal microbiota is driven by rapid uptake and conversion of simple carbohydrates. The ISME journal, 2012. 6(7): p. 1415.

33. Chen, Y, et al., Dysbiosis of small intestinal microbiota in liver cirrhosis and its association with etiology. Scientific reports, 2016. 6: p. 34055.

34. Al Mardini, H., K. Bartlett, and C.O. Record, Blood and brain concentrations of mercaptans in hepatic and methanethiol induced coma. Gut, 1984. 25(3): p. 284–290.

35. Contreras, A., et al., Importance of Dialister pneumosintes in human periodontitis. Oral microbiology and immunology, 2000. 15(4): p. 269–272.

36. Moore, W.E.C. and L.V.H. Moore, The bacteria of periodontal diseases. Periodontology 2000, 1994. 5(1): p. 66–77.

37. Coker, O.O., et al., Mucosal microbiome dysbiosis in gastric carcinogenesis. Gut, 2018. 67(6): p. 1024–1032.

38. Gomez, C.A., et al., First case of infectious endocarditis caused by Parvimonas micra. Anaerobe, 2015. 36: p. 53–55.

39. Baghban, A. and S. Gupta, Parvimonas micra: a rare cause of native joint septic arthritis. Anaerobe, 2016. 39: p. 26–27.

40. Uemura, H., et al., Parvimonas micra as a causative organism of spondylodiscitis: a report of two cases and a literature review. International Journal of Infectious Diseases, 2014. 23: p. 53–55.

41. Perry, A. and P. Lambert, Propionibacterium acnes: infection beyond the skin. Expert review of anti-infective therapy, 2011. 9(12): p. 1149–1156.

42. Douglas, H.C. and S.E. Gunter, The taxonomic position of Corynebacterium acnes. Journal of bacteriology, 1946. 52(1): p. 15.

43. Ohigashi, S., et al., Changes of the intestinal microbiota, short chain fatty acids, and fecal pH in patients with colorectal cancer. Digestive diseases and sciences, 2013. 58(6): p. 1717–1726.

44. Trapnell, C., et al., Transcript assembly and quantification by RNA-Seq reveals unannotated transcripts and isoform switching during cell differentiation. Nature biotechnology, 2010. 28(5): p. 511.

45. Patro, R., S.M. Mount, and C. Kingsford, Sailfish enables alignment-free isoform quantification from RNA-seq reads using lightweight algorithms. Nature biotechnology, 2014. 32(5): p. 462.

46. Xia, L.C., et al., Accurate genome relative abundance estimation based on shotgun metagenomic reads. PloS one, 2011. 6(12): p. e27992.

47. Li, D., et al., MEGAHIT: an ultra-fast single-node solution for large and complex metagenomics assembly via succinct de Bruijn graph. Bioinformatics, 2015. 31(10): p. 1674–1676.

48. Kang, D.D., et al., MetaBAT, an efficient tool for accurately reconstructing single genomes from complex microbial communities. PeerJ, 2015. 3: p. e1165.

49. Meyer, F., et al., AMBER: Assessment of Metagenome BinnERs. GigaScience, 2018.

50. Li, H. and R. Durbin, Fast and accurate short read alignment with Burrows–Wheeler transform. bioinformatics, 2009. 25(14): p. 1754–1760.

51. Storey, J.D., The positive false discovery rate: a Bayesian interpretation and the q-value. The Annals of Statistics, 2003. 31(6): p. 2013–2035.

